# Genomic selection for accelerated heartwood formation in Pedunculate oak (*Quercus robur* L.) using whole-genome sequencing

**DOI:** 10.1101/2025.07.22.666121

**Authors:** Albin Lobo, James Doonan, Jon Kehlet Hansen, Chatchai Kosawang, Jing Xu, Jill K Olofsson, Erik Dahl Kjær

## Abstract

Heartwood traits in trees are critical for timber quality but are notoriously difficult to phenotype due to their late expression and the need for destructive sampling. In this proof-of-concept study, we demonstrate that combining genome-wide association studies (GWAS) with Bayesian genomic prediction models provides an effective strategy to overcome these challenges. By using GWAS to preselect trait associated SNPs and integrating them into predictive models, we substantially improve the accuracy of genomic predictions for heartwood related traits in oaks. Our approach allows for reliable selection of superior genotypes at the seedling stage, long before heartwood traits can be directly measured, thus enabling early and cost-effective breeding decisions. We also identify the number of rings in sapwood as a genetically controlled, easily measured proxy trait that enhances selection strategies for heartwood content. Together, these findings provide a scalable framework for integrating genomics into operational tree breeding programs and demonstrate how combining GWAS and genomic prediction can accelerate the improvement of complex wood traits in long lived forest tree species.

## Introduction

Due to their long generation time, tree breeding programs initiated as early as the mid-1950s are still in their second or third generation (Koskela *et al*., 2014). Significant progress has been made in understanding the molecular mechanisms of breeding using microsatellites and single nucleotide polymorphism (SNP) data. However, many forest tree species still lag behind crop plants in terms of genomic resources. This disparity is largely due to their complex genomes, often characterized by high levels of heterozygosity and repetitive DNA as well as long generation times and large, outcrossing populations. Although molecular marker-based approaches have been explored in tree breeding, their success has been limited (Holliday *et al*., 2017), as they either explain only a small percentage of phenotypic variation or are ineffective in studying polygenic traits (Isik, 2014).

Genomic selection (GS) is increasingly recognized as a powerful tool for accelerating genetic improvement in long-lived organisms like trees (Isik, 2014; El-Kassaby *et al*., 2024). Recent advancements in genomics and phenomics enable the decoding of entire genomes and precise phenotyping of economically significant complex traits in trees such as wood quality. GS integrates quantitative genetic prediction models with SNP markers, facilitating the identification of individuals with more favorable alleles. With the advent of full genome sequences for tree species, GS has been successfully applied, yielding significant genetic gains with high accuracy in different tree genera such as *Pinus* (Resende *et al*., 2012a) and *Eucalyptus* (Resende *et al*., 2012c) and *Picea* (Beaulieu *et al*., 2020; Lenz *et al*., 2020). Key genomic resources, such as the oak genome sequence (Plomion *et al*., 2016; ‘Quercus robur genome assembly dhQueRobu3.1’), are now available, enabling the selection of elite genotypes based on genomic data.

*Quercus robur*, commonly known as the European or pedunculate oak (hereafter referred to as oak), is a key species in European forestry. The species can grow on a broad range of sites and is important for forest biodiversity. The timber is a high value product due to strength, durability, and versatility, and the species is therefore of significant economic importance in many countries (Eaton *et al*., 2016). However, its long rotation period poses a significant economic challenge. Thinning operations occur early in the plantation’s lifecycle and ensure a steady supply of wood, but they mostly yield sapwood-rich timber, which has limited value and is primarily used for pulp and energy purposes. Increasing the proportion of heartwood at an earlier stage could allow the production of high-quality timber even during thinning, thereby boosting revenue from these intermediate harvests and enhancing the overall yield of valuable wood products.

Genetic improvement of wood quality has played a crucial role in forest tree breeding for enhancing economic returns (Jansson *et al*., 2017), although breeding efforts specifically for wood quality in oak have been limited. However, genetic variation in sapwood percentage has been reported in the species (Savill *et al*., 1993; Sohar *et al*., 2012), highlighting the potential for breeding this trait. Traditionally, tree improvement programs have focused on selecting traits such as height, diameter, and stem form as indicators of wood quality, because phenotyping of heartwood formation is relatively time consuming. This stresses the importance of making selections for heartwood formation as reliably as possible, and one approach here is to use GS. Understanding the genetic control underlying heartwood formation is highly valuable in order to guide early selection for this trait. Early selection for superior heartwood content ensures that the simultaneous selection of other key traits, such as growth rate, is not compromised. Therefore, there is a pressing need to develop faster and more cost-effective breeding tools and strategies to reliably select for superior heartwood content.

The objectives of this study were to evaluate the potential of GS for early heartwood formation in oaks by identifying informative SNP markers and developing predictive models for heartwood related traits. Specifically, we aimed to (1) quantify genetic variation and heritability of heartwood traits, (2) implement genome-wide association studies (GWAS) to select the most informative SNPs associated with heartwood formation, and (3) use these trait-associated SNPs to construct Bayesian genomic prediction models to assess the predictive power of selected markers. By integrating GWAS with GS, we aim to demonstrate a ‘proof of concept’ for accelerating the identification of superior genotypes, facilitating early selection in oak breeding programs. This approach provides a foundation for the future development of customized, cost-effective genotyping platforms and supports the incorporation of heartwood content as a key trait in oak improvement strategies for European forestry.

## Materials and Methods

### Study material and phenotyping

The study was conducted in an oak progeny trial (common garden) in Vejbæk in southern Jutland, Denmark (54.88°N, 9.30°E). The trial was established in 2002 with a randomized, unbalanced block design with trees planted at a spacing of 3 × 1 meters. For this study, three trees from each of 100 open-pollinated families (300 trees in total) were selected.

Wood cores were collected from the 300 trees at a height of 50 cm above ground level using a 5-mm increment borer. After extraction, the cores were air-dried and securely mounted on core holders with a water-soluble wooden adhesive. They were then sanded to create a smooth, flat surface, and phenotypic traits, including heartwood width, heartwood area, sapwood width, and the number of rings in the sapwood were measured using the Velmex Tree Ring Measuring System (Velmex Inc., USA). Heartwood area was estimated from heartwood width by assuming a circular cross-sectional area.

### Whole genome sequencing, SNP calling and GWAS

DNA was extracted from leaf tissue of 300 oak trees using a Nucleospin Plant II kit (Macherey-Nagel, Germany) following the manufacturer’s instructions. DNA quantitation was done using a Nanodrop 2000 spectrophotometer (ThermoFisher Scientific, USA) and a QUBIT fluorometer 2.0 (ThermoFisher Scientific, USA) prior to whole genome sequencing with the Illumina NovaSeq 6000 in 150PE mode at IGATech (Udine, Italy). Capturing probes had been previously designed (Lesur *et al*., 2018) and were additionally designed from the newly available oak genome (‘Quercus robur genome assembly dhQueRobu3.1’).

To develop and evaluate genomic prediction models for heartwood-related traits, we performed GWAS using phenotypic observations from all individuals. This approach allowed us to preselect the top 50000 SNPs most significantly associated with each trait, thereby reducing dimensionality while retaining biologically informative markers. As demonstrated by El-Kassaby *et al*., 2024, leveraging trait-associated SNPs as input to genomic selection models can enhance predictive performance by enriching the marker set for loci with large or moderate effect sizes. By filtering the SNP dataset based on GWAS results, we aimed to capture a greater proportion of the additive genetic variance relevant to each trait, while minimizing noise from uninformative or redundant markers. To benchmark the utility of GWAS-based SNP selection, we implemented a comparative genomic prediction analysis using a second, randomly selected set of 50,000 SNPs drawn from the full SNP dataset. These random SNPs were matched in number and quality filters (e.g., minor allele frequency, call rate) but were not prioritized based on trait associations. This allowed us to test the hypothesis that prediction accuracy is significantly higher when using trait-associated SNPs compared to random ones.

GWAS was performed using a univariate linear mixed model within the GEMMA v0.94.1 software toolkit(Zhou & Stephens, 2012). Single nucleotide polymorphisms (SNPs) were called at variable sites first using BCFtools v1.16 (Li, 2011), removing those with a quality score below 20 (-q 20) and filtering for low quality (LowQual) to create a file in variant call format (VCF). Subsequently, the VCF file was converted to BIMBAM format using Plink2 v1.90beta6.24 (Purcell *et al*., 2007). Using BIMBAM format files, a centred relationship matrix was estimated using GEMMA v0.94.1. Genome wide associations incorporated the relationship matrix and used a minimum allele frequency (MAF) of 0.01 to exclude rare variants. Subsequent results were ranked according to each SNP’s likelihood ratio test statistic (p_lrt). Resultant quantile-quantile (QQ) and Manhattan plots were drawn using the R package qqman v0.1.9 (Turner, 2018). Top ranked SNPs were converted into BLUP format using the Angsd v0.935 software toolkit (Korneliussen *et al*., 2014).

### Quantitative genetic analysis of the phenotypes

We predicted the genomic estimated breeding values (GEBVs) of individuals based on their genome-wide single nucleotide polymorphism (SNP) profiles. This approach enables the identification and selection of genotypes with superior phenotypic performance for heartwood-related traits.

A genomic best linear unbiased prediction (GBLUP) model was implemented, where additive genetic relationships among individuals were inferred from genome-wide SNP markers. The marker data were used to construct a genomic relationship matrix (GRM), which captures the average genomic similarity among individuals. This matrix replaces the traditional pedigree-based relationship matrix, allowing more accurate estimation of breeding values by integrating realized genomic relationships (Visscher *et al*., 2008; Hayes *et al*., 2009; Grattapaglia & Resende, 2011; Resende *et al*., 2012b).

The GBLUP model was specified as:

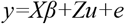

where *y* is the vector of phenotypic observations for heartwood traits, *X* and *β* represent the design matrix and vector of fixed effect respectively, *Z* is the incidence matrix linking observations to genomic effects, *u* is the vector of additive genetic effects assumed to follow u=σ^2^G_A_), where G_A_ is the GRM and *e* is the residual error term.

Heritability estimates of the phenotypic traits were calculated using the genomic relationship matrix (G-matrix) approach. The G-matrix was constructed based on genome-wide SNP markers obtained from whole-genome sequencing. A restricted maximum likelihood (REML) method was applied using a linear mixed model to partition phenotypic variance into genetic and residual components. The narrow-sense heritability (*h*^*2*^) was estimated as the ratio of additive genetic variance (*σ*^*2*^_*A*_) to total phenotypic variance (*σ*^*2*^_*P*_):

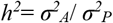

All statistical analyses were performed using the ASReml-W v4.2 software. Model fit and significance of variance components were evaluated via likelihood ratio tests.

### Bayesian genomic selection model

To evaluate the potential of genomic prediction for heartwood-related traits in oaks, we implemented a Bayesian genomic selection model using the R package BGLR (Pérez & de los Campos, 2014). Genotypic data (50,000 most significantly associated SNPs for each trait) were formatted into a matrix where rows represented individual trees and columns represented SNP markers. The phenotypic data were matched and filtered to retain only those individuals present in both datasets. All data processing and model fitting were conducted in R version 4.3.2

We used the BayesB model to account for variable selection and shrinkage of SNP effects (Isik *et al*., 2017). The model was fitted using 15,000 Markov Chain Monte Carlo (MCMC) iterations, with a burn-in of 3,000 iterations. The model equation was:

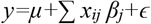

where *y* is the vector of phenotypes for the trait of interest, *μ* is the overall mean, *x*_*ij*_ is the genotype at SNP *j* for individual *i, β*_*j*_ is the marker effect for SNP *j*, and *ϵ* is the residual error.

To evaluate model performance, we performed 5-fold cross-validation on the full set of 300 trees. Each run involved partitioning the data into five equal subsets; four subsets (80%) were used for model training, and one subset (20%) for validation. This was repeated 10 times with different random splits, yielding 50 estimates of prediction accuracy. Accuracy was computed as the Pearson correlation between predicted genetic values and observed phenotypes (Isik *et al*., 2017). The process was applied separately to models using GWAS-selected SNPs and randomly selected SNPs. This comparative approach allowed us to quantify the added value of trait-associated SNPs over randomly chosen markers. A higher prediction accuracy with GWAS-derived SNPs would support their utility in marker-based selection and provide a stronger rationale for developing cost-effective, trait-informed genotyping platforms in oak breeding programs.

All relevant model outputs, including predicted values, observed values, and accuracy metrics, were exported for each trait. A summary table was prepared showing mean prediction accuracy with standard deviation (SD) for all phenotypic traits based on their respective top 50,000 SNPs (Table 2).

## Results

### Quantitative genetic variation in heartwood content in oak

Table 1 summarizes the estimates of additive genetic variance (*V*_*a*_), residual variance (*V*_*e*_), narrow-sense heritability (*h*^*2*^), and associated standard errors for six phenotypic traits assessed. *V*_*a*_ was significant for all the traits analyzed. Heritability estimates ranged from 0.47 for diameter and sapwood width to 0.72 for heartwood area, indicating moderate to high additive genetic control. Heartwood area showed the highest heritability (0.72), suggesting strong potential for genetic improvement through selection. All traits exhibited similar standard errors (0.29–0.32), reflecting consistent estimation precision across traits.

**Table 1:** Additive variance (*V*_*a*_), Environmental variance (*V*_*e*_) and Narrow sense heritability (*h*^*2*^) estimate along with its standard error (*SE*) of the phenotypic traits.

| Trait | $V_a$ | $V_e$ | $h^2$ | Std.error |
| --- | --- | --- | --- | --- |
| Diameter (mm) | 1065.69 | 1210.47 | 0.47 | 0.30 |
| Heartwood Width (mm) | 180.22 | 182.16 | 0.50 | 0.31 |
| Heartwood Area (mm<sup>2</sup>) | 3.15E+07 | 1.22E+07 | 0.72 | 0.32 |
| Sapwood Width (mm) | 45.62 | 51.05 | 0.47 | 0.29 |
| No.of rings in Sapwood | 2.55 | 2.53 | 0.50 | 0.30 |

**Table 2:** Prediction accuracy (mean ± SD) for heartwood-related traits using SNPs preselected by GWAS compared to accuracy using a random SNP set.

| Trait | Prediction Accuracy (mean ± SD)<br>using SNPs preselected through GWAS | Prediction Accuracy (mean ± SD)<br>using random SNPs |
| --- | --- | --- |
| Diameter (mm) | 0.93 ± 0.02 | 0.20 ± 0.15 |
| Heartwood Width (mm) | 0.94 ± 0.02 | 0.17 ± 0.15 |
| Heartwood Area (mm<sup>2</sup>) | 0.93 ± 0.02 | 0.17 ± 0.14 |
| Sapwood Width (mm) | 0.94 ± 0.02 | 0.16 ± 0.13 |
| No.of rings in Sapwood | 0.95 ± 0.02 | 0.13 ± 0.14 |

### Preselection of informative SNPs using GWAS

Variant calling from whole genome resequencing identified 45.5 million genome-wide single nucleotide polymorphisms (SNPs). Phenotypic trait associations were conducted using the complete SNP dataset, and resultant *P* values were visualized in Manhattan plots (Figure1). SNP sites under linkage disequilibrium can be identified as peaks or towers of clustered SNPs within neighboring genomic regions. The significance value of each SNP is shown in relation to the standard GWAS significance threshold of 5×10^−8^, which was included in the Manhattan plots for illustrative purposes but was not used to further interpret results. The distribution and significance of variable SNP sites in Figure 1 reveal the genetic architecture of associated traits. The top 50,000 SNPs based on *P* value significance were used as input for genomic selection analysis.

**Figure 1.**
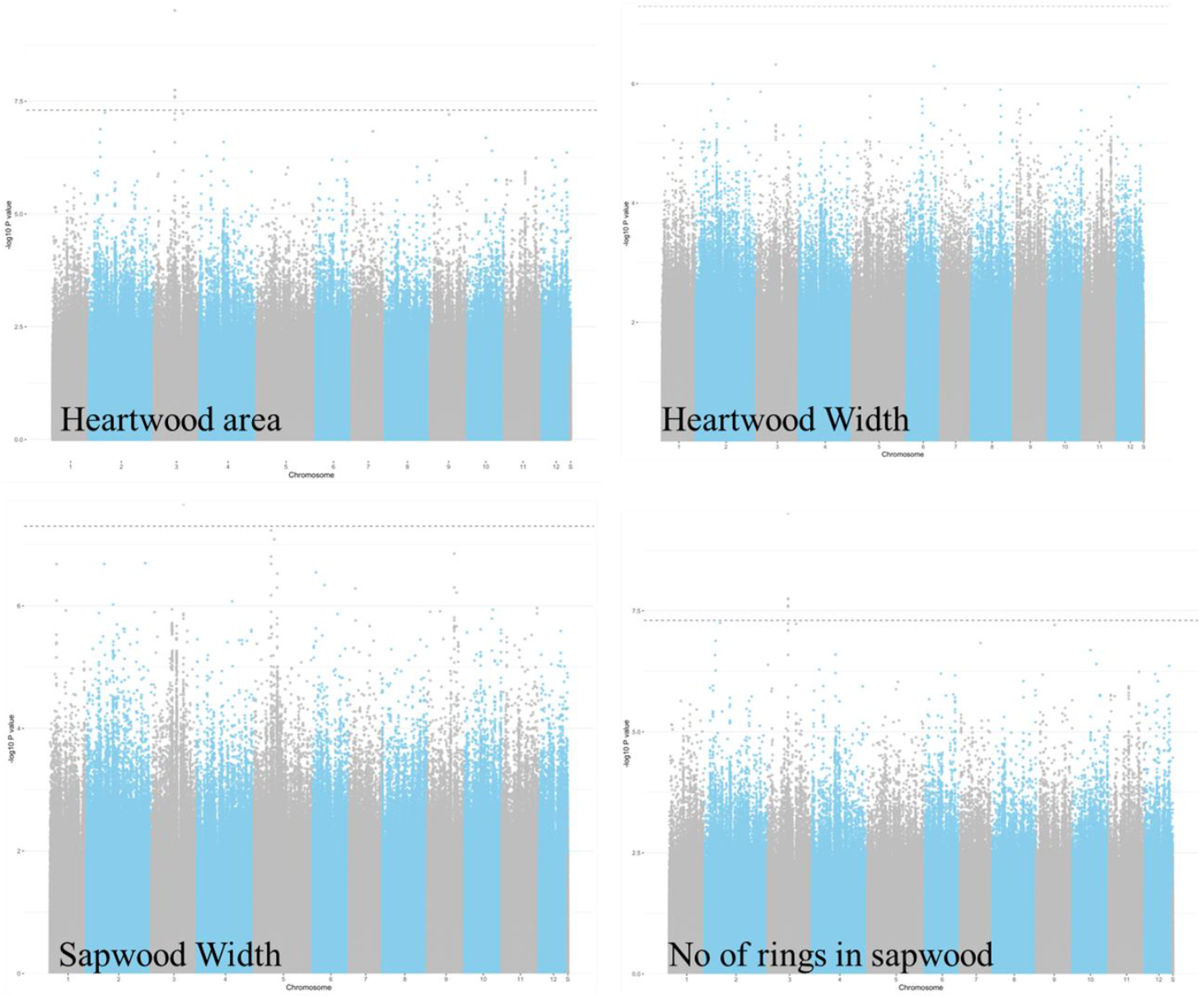
Manhattan plots of genome-wide association results for the four heartwood related traits: heartwood width, heartwood area, sapwood width, and number of rings in sapwood.

### Genomic prediction of heartwood traits using Bayesian models

The individuals with the highest GEBVs were considered as candidates for selection to enhance heartwood content in future breeding cycles. Table 2 summarizes the predictive ability (Pearson correlation between observed and predicted values) for eight phenotypic related traits in oaks studied. For each trait, only the top 50,000 most associated SNPs (see GWAS results above) were included in the model. Values represent mean ± standard deviation of the correlation coefficients across replicates.

## Discussion

### High genetic control of heartwood traits facilitates targeted breeding and high throughput phenotyping

The heritability estimates for general growth traits in oak trees suggest a moderate to high degree of genetic control, underscoring the potential for accelerating breeding progress.

Narrow-sense heritability for diameter was moderately high (*h*^*2*^ = 0.47), which points to a strong additive genetic component and relatively low environmental variance for these traits in our dataset. Our findings are consistent with previous studies reporting moderate to high heritability for growth traits in forest trees such as *Eucalyptus* (Resende *et al*., 2012c) and *Pinus taeda* (Resende *et al*., 2012b), confirming the general trend of strong additive genetic control in forest tree growth. While standard errors for heritability estimates were relatively large (~0.3), likely due to sample size, the overall pattern suggests a high breeding potential for the traits.

Heartwood related traits were also found to be under strong genetic control. The heartwood area showed a high heritability estimate (*h*^*2*^=0.72), indicating that genomic selection for this trait could lead to significant improvements in heartwood yield. Other traits related to heartwood formation, such as heartwood width (*h*^*2*^=0.50) and sapwood width (*h*^*2*^=0.47), also demonstrated moderately high heritability, consistent with previous findings that wood quality traits in oaks can be genetically controlled (Mosedale *et al*., 1996).

The high heritability observed for the number of rings in the sapwood (*h*^*2*^ = 0.50) underscores the strong genetic control of this trait and reinforces its potential utility in genomic selection strategies targeting overall tree growth and cambial age-related characteristics. This finding is particularly valuable as it highlights the number of sapwood rings as a practical, rapid, and cost-effective phenotyping proxy for selecting trees with superior heartwood content. Since the number of sapwood rings is under genetic control, future breeding programs could rely on minimally invasive sampling techniques. Specifically, a small increment core from the outer sapwood region would suffice to count the number of sapwood rings. This simple measurement could then serve as an indirect but reliable criterion for selecting individuals with enhanced heartwood development, thereby streamlining the selection process and accelerating genetic improvement.

### Genomic selection enables accurate early prediction of heartwood traits in oaks, facilitating selection at the seedling stage for hard-to-phenotype characteristics

For the number of rings in sapwood trait, the prediction accuracy was high, with a mean ± standard deviation of 0.95 ± 0.02, indicating strong model performance and consistency across replicates. The cross-validation strategy (refer to materials and methods) provides a realistic assessment of the model’s ability to generalize to new individuals and reflects its practical utility in breeding programs. In operational settings, genomic selection allows early selection of superior individuals by genotyping seedlings that have not yet been phenotyped (Jannink *et al*., 2010). For traits such as the number of rings in sapwood, which typically require destructive sampling or many years of growth to measure, genomic prediction enables selection at the seedling stage. This eliminates the need to wait for mature trees to express the trait, thereby reducing costs and enabling earlier selection decisions. The high prediction accuracy observed here suggests that this approach is highly effective for this trait and can be confidently applied to support early selection in oak breeding programs.

The ability to achieve high predictive accuracy for heartwood traits analyzed in this study, which are difficult or impossible to measure in young trees or in living individuals without destructive sampling, opens up new possibilities for early selection in breeding programs. By leveraging genomic selection with trait-informed marker sets, it becomes feasible to make informed breeding decisions at the seedling stage, thereby shortening breeding cycles and improving genetic gain for commercially and ecologically important wood properties.

### Genomic prediction of heartwood traits benefits from GWAS informed SNP selection and Bayesian modeling

Our study demonstrates that integrating genome-wide association studies (GWAS) to preselect informative SNPs with Bayesian genomic prediction models provides a powerful framework for predicting complex, hard-to-phenotype traits such as heartwood characteristics in oaks. Specifically, we achieved a high prediction accuracy for heartwood area (r = 0.93; Table 2) when using SNPs preselected based on GWAS results and applying a Bayesian model for genomic prediction. In stark contrast, when an equivalent number of SNPs were selected at random, prediction accuracy dropped significantly to 0.17 (Table 2), underscoring the importance of informed marker selection.

These findings highlight two key points. First, the use of GWAS enables the identification of trait-associated genomic regions that are likely to capture relevant additive genetic variance, even for complex traits with potentially polygenic architecture. Second, Bayesian models, particularly those that accommodate variable selection and shrinkage (e.g., BayesB), are well suited for incorporating this prior information to generate accurate genomic predictions. This approach represents a significant advancement in the application of genomic tools for forest tree improvement, where long generation times, complex phenotyping, and late trait expression have traditionally hindered selection efficiency. Our results provide a strong argument for integrating GWAS informed marker selection with Bayesian genomic prediction as a standard strategy for improving traits that are otherwise challenging to assess directly.

The findings of this study demonstrate the feasibility of applying genomic prediction to accelerate breeding for complex wood traits such as heartwood area in oak. With increasing availability of high-resolution genomic data, the integration of trait-associated SNPs into predictive models offers a powerful tool to reduce breeding cycle time and enhance selection efficiency. In practical terms, the approach we employed, combining GWAS based marker selection with Bayesian genomic prediction, lays the foundation for a streamlined, genomics-enabled tree improvement pipeline. After training and validating the model using a subset of phenotyped and genotyped individuals, the resulting model can be applied to predict the genetic merit of unphenotyped trees based solely on their genomic profiles (Nadeau *et al*., 2023). This enables early and cost-effective selection of superior individuals prior to trait expression, which is especially valuable for traits like heartwood that manifest only in mature trees.

In a breeding context, the implementation of this approach would involve genotyping candidate seedlings or clonal material using the preselected SNP panel and predicting their breeding values using the validated genomic prediction model. Individuals with the highest predicted genetic merit could then be prioritized for deployment in improved plantations, inclusion in clonal seed orchards, or as parents in the next breeding cycle. By doing so, selection decisions can be made in the seedling stage, bypassing the need for multi-year phenotypic evaluations. Importantly, this genomic selection strategy can be continuously refined. As more phenotypic and genotypic data become available, prediction models can be updated to improve their accuracy and generalizability across environments or genetic backgrounds. The iterative nature of this approach aligns well with long-term tree improvement programs and has the potential to significantly accelerate genetic gain.

In conclusion, our results validate the predictive power of genomic models and demonstrate their practical value for operational breeding programs targeting traits that are difficult to accurately phenotype, time consuming, or costly to measure. This transition from phenotype-based to genomics-supported selection marks a significant step forward in developing resource efficient forest genetic improvement strategies. The strong association between the number of rings in sapwood and heartwood content, offers options for developing a cost effective, high throughput phenotyping proxy for heartwood traits in mature trees. The tools will support rapid breeding for increased heartwood content.

## Data availability

Whole genome sequences are available on the GenBank/EBI/DDBJ under BioProject accession PRJDB35455.

## Acknowledgement

We gratefully acknowledge financial support from the Innovation Fund Denmark for funding the FASTWOOD project (0139-00011B). We extend our sincere thanks to our collaborators at the Institute of Forest Genetics and Tree Breeding in India for their valuable partnership, and to the Danish Nature Agency for granting access to the field trial site. We also thank Lars Nørgaard Hansen and Carsten Tom Nørgaard for their assistance during fieldwork, and Martina Stoop for measuring heartwood. Finally, we appreciate Lisbeth Garbrecht Thygesen and Andrea Ponzecchi for their helpful comments on the manuscript.

## References

Beaulieu J, Nadeau S, Ding C, Celedon JM, Azaiez A, Ritland C, Laverdière J-P, Deslauriers M, Adams G, Fullarton M, et al. 2020. Genomic selection for resistance to spruce budworm in white spruce and relationships with growth and wood quality traits. Evolutionary Applications 13: 2704–2722.

Eaton E, Caudullo G, Oliveira S, De Rigo D. 2016. Quercus robur and Quercus petraea in Europe: distribution, habitat, usage and threats. European atlas of forest tree species: 160– 163.

El-Kassaby YA, Cappa EP, Chen C, Ratcliffe B, Porth IM. 2024. Efficient genomics-based ‘end-to-end’ selective tree breeding framework. Heredity 132: 98–105.

Grattapaglia D, Resende MDV. 2011. Genomic selection in forest tree breeding. Tree Genetics & Genomes 7: 241–255.

Hayes BJ, Visscher PM, Goddard ME. 2009. Increased accuracy of artificial selection by using the realized relationship matrix. Genetics Research 91: 47–60.

Holliday JA, Aitken SN, Cooke JEK, Fady B, González-Martínez SC, Heuertz M, Jaramillo-Correa J, Lexer C, Staton M, Whetten RW, et al. 2017. Advances in ecological genomics in forest trees and applications to genetic resources conservation and breeding. Molecular Ecology 26: 706–717.

Isik F. 2014. Genomic selection in forest tree breeding: the concept and an outlook to the future. New Forests 45: 379–401.

Isik F, Holland J, Maltecca C. 2017. Genomic Selection. In: Isik F, Holland J, Maltecca C, eds. Genetic Data Analysis for Plant and Animal Breeding. Cham: Springer International Publishing, 355–384.

Jannink J-L, Lorenz AJ, Iwata H. 2010. Genomic selection in plant breeding: from theory to practice. Briefings in Functional Genomics 9: 166–177.

Jansson G, Hansen JK, Haapanen M, Kvaalen H, Steffenrem A. 2017. The genetic and economic gains from forest tree breeding programmes in Scandinavia and Finland. Scandinavian Journal of Forest Research 32: 273–286.

Korneliussen TS, Albrechtsen A, Nielsen R. 2014. ANGSD: Analysis of Next Generation Sequencing Data. BMC Bioinformatics 15: 356.

Koskela J, Vinceti B, Dvorak W, Bush D, Dawson IK, Loo J, Kjaer ED, Navarro C, Padolina C, Bordács S, et al. 2014. Utilization and transfer of forest genetic resources: A global review. Forest Ecology and Management 333: 22–34.

Lenz PRN, Nadeau S, Mottet M-J, Perron M, Isabel N, Beaulieu J, Bousquet J. 2020. Multi-trait genomic selection for weevil resistance, growth, and wood quality in Norway spruce. Evolutionary Applications 13: 76–94.

Lesur I, Alexandre H, Boury C, Chancerel E, Plomion C, Kremer A. 2018. Development of Target Sequence Capture and Estimation of Genomic Relatedness in a Mixed Oak Stand. Frontiers in Plant Science 9.

Li H. 2011. A statistical framework for SNP calling, mutation discovery, association mapping and population genetical parameter estimation from sequencing data. Bioinformatics 27: 2987–2993.

Mosedale JR, Charrier B, Janin G. 1996. Genetic control of wood colour, density and heartwood ellagitannin concentration in European oak (Quercus petraea and Q. robur). Forestry: An International Journal of Forest Research 69: 111–124.

Nadeau S, Beaulieu J, Gezan SA, Perron M, Bousquet J, Lenz PRN. 2023. Increasing genomic prediction accuracy for unphenotyped full-sib families by modeling additive and dominance effects with large datasets in white spruce. Frontiers in Plant Science 14.

Pérez P, de los Campos G. 2014. Genome-Wide Regression and Prediction with the BGLR Statistical Package. Genetics 198: 483–495.

Plomion C, Aury J, Amselem J, Alaeitabar T, Barbe V, Belser C, Bergès H, Bodénès C, Boudet N, Boury C, et al. 2016. Decoding the oak genome: public release of sequence data, assembly, annotation and publication strategies. Molecular Ecology Resources 16: 254–265.

Purcell S, Neale B, Todd-Brown K, Thomas L, Ferreira MAR, Bender D, Maller J, Sklar P, Bakker PIW de, Daly MJ, et al. 2007. PLINK: A Tool Set for Whole-Genome Association and Population-Based Linkage Analyses. The American Journal of Human Genetics 81: 559–575.

Quercus robur genome assembly dhQueRobu3.1. NCBI.

Resende MFR, Muñoz P, Acosta JJ, Peter GF, Davis JM, Grattapaglia D, Resende MDV, Kirst M. 2012a. Accelerating the domestication of trees using genomic selection: accuracy of prediction models across ages and environments. New Phytologist 193: 617–624.

Resende MFR Jr, Muñoz P, Resende MDV, Garrick DJ, Fernando RL, Davis JM, Jokela EJ, Martin TA, Peter GF, Kirst M. 2012b. Accuracy of Genomic Selection Methods in a Standard Data Set of Loblolly Pine (Pinus taeda L.). Genetics 190: 1503–1510.

Resende MDV, Resende MFR, Sansaloni CP, Petroli CD, Missiaggia AA, Aguiar AM, Abad JM, Takahashi EK, Rosado AM, Faria DA, et al. 2012c. Genomic selection for growth and wood quality in Eucalyptus: capturing the missing heritability and accelerating breeding for complex traits in forest trees. New Phytologist 194: 116–128.

Savill PS, Kanowski PJ, Gourlay, ID, Jarvis AR. 1993. Genetic and intratree variation in the number of Sapwood Rings in Quercus robur and Q. petraea. Silvae genetica 42: 371–375.

Sohar K, Vitas A, Läänelaid A. 2012. Sapwood estimates of pedunculate oak (Quercus robur L.) in eastern Baltic. Dendrochronologia 30: 49–56.

Turner SD. 2018. qqman: an R package for visualizing GWAS results using Q-Q and manhattan plots. Journal of Open Source Software 3: 731.

Visscher PM, Hill WG, Wray NR. 2008. Heritability in the genomics era — concepts and misconceptions. Nature Reviews Genetics 9: 255–266.

Zhou X, Stephens M. 2012. Genome-wide efficient mixed-model analysis for association studies. Nature Genetics 44: 821–824.

